# Integrative analysis reveals unique features of the Smc5/6 complex

**DOI:** 10.1101/2020.12.31.424863

**Authors:** You Yu, Shibai Li, Zheng Ser, Tanmoy Sanyal, Koyi Choi, Bingbing Wan, Andrej Sali, Alex Kentsis, Dinshaw J. Patel, Xiaolan Zhao

## Abstract

Structural maintenance of chromosomes (SMC) complexes are critical chromatin modulators. In eukaryotes, the cohesin and condensin SMC complexes organize chromatin, while the Smc5/6 complex directly regulates DNA replication and repair. The molecular basis for Smc5/6’s distinct functions is currently poorly understood. Here, we report an integrative structural study of the budding yeast Smc5/6 complex using electron microscopy, cross-linking mass spectrometry, and computational modeling. We show that while the complex shares a similar overall organization with other SMC complexes, it possesses several unique features. In contrast to the reported folded-arm structures of cohesin and condensin, our data suggest that Smc5 and Smc6 arm regions do not fold back on themselves. Instead, these long filamentous regions interact with subunits uniquely acquired by the Smc5/6 complex, namely the Nse2 SUMO ligase and the Nse5-Nse6 subcomplex. Further, we show that Nse5-Nse6 subcomplex adopts a novel structure with an extensive dimerization interface and multiple domains contacting other subunits of the Smc5/6 complex. We also provide evidence that the Nse5-Nse6 module uses its SUMO-binding motifs to contribute to Nse2-mediated sumoylation. Collectively, our integrative multi-scale study identifies distinct structural features of the Smc5/6 complex and functional cooperation amongst its co-evolved unique subunits.

## Introduction

Structural maintenance of chromosomes (SMC) complexes regulate genome organization and maintenance in both prokaryotic and eukaryotic cells. Each complex contains a pair of SMC subunits and a set of non-SMC subunits (1). Studies of several SMC proteins reveal that they form tripartite filamentous structures. An SMC subunit folds back on itself at its middle ‘hinge’ region, enabling its N- and C-terminal ATPase domains to associate forming a ‘head’ region, and its two long coiled-coil regions located in-between the hinge and the head to pair in an anti-parallel manner, forming an ‘arm’ region (Fig. S1A) (1). The two SMC subunits of each complex form its backbone and can associate with each other at hinge, head and arm regions (1).

Much of the molecular understanding of SMC complexes has come from studies of those acting as DNA organization and separation factors, such as prokaryotic Smc-ScpAB and MukBEF complexes and eukaryotic cohesin and condensin. These complexes can entrap and loop DNA, resulting in DNA tethering and folding (2–4). One emerging feature of these complexes is that their long arm regions bend sharply at so-called ‘elbow’ sites (Fig. S1A). Elbow bending causes the hinge to contact the head-proximal coiled-coil or head-bound non-SMC proteins, a conformation thought to facilitate DNA loop extrusion (5–8). In these SMCs, the head and hinge regions that associate with other proteins and/or DNA are conserved, while the arm regions are not and act mainly as connecting elements (9).

Differing from cohesin (containing Smc1/3) and condensin (containing Smc2/4), the third eukaryotic SMC complex, containing Smc5 and Smc6, does not appear to affect chromatid intertwining or mitotic chromosome structures (10–12). Rather, the Smc5/6 complex directly regulates DNA replication and recombinational repair (13–15). These unique functions correlate with the acquisition of a special set of six subunits, namely <u>N</u>on-<u>S</u>MC <u>E</u>lements (Nse)1-6. Three Nse subunits, Nse2, Nse5 and Nse6, are not found in any other SMC complexes in either prokaryotes or eukaryotes. Nse2 (aka Mms21) is a SUMO ligase that promotes the sumoylation of more than a dozen genome maintenance factors (16–19). Nse5 and 6 are thought to act distinctly from Nse2 by forming a subcomplex that recruits the Smc5/6 complex to DNA damage sites (13-15, 20–22). The acquisition of the Nse2, Nse5 and Nse6 subunits is one of the most unique features that sets the Smc5/6 complex apart from other SMC complexes.

Our understanding of how the Smc5/6 complex gained unique functions among the SMC family of complexes is hindered by the limited structural information of its holo-complex. Studies of subunits and their fragments or subcomplexes have provided insights into potential intersubunit interactions (23–25). However, these data may not reflect structures and interactions within the entire complex. Thus, it is imperative to determine, in the context of the holo-complex, whether Smc5 and Smc6 adopt distinct conformations relative to other SMCs, how they associate with Nse subunits, and what the functional relationships are amongst the complex-specific Nse2, 5 and 6 subunits. Here, we provide an integrative structural analysis of the Smc5/6 holo-complex isolated from budding yeast that addresses these challenges. We use negative staining electron microscopy, cross-linking mass spectrometry (CL-MS), single particle cryo-EM, structural modeling and functional analyses to identify several unique features of the Smc5/6 complex that distinguish it from the other SMC complexes. We also provide evidence that the co-evolved Nse2, Nse5, and Nse6 subunits are connected at both structural and functional levels.

## Results

### The overall structure of the Smc5/6 holo-complex

To gain structural insight into the Smc5/6 holo-complex, we co-expressed all eight subunits of the budding yeast complex in its native host cells (see Methods). The holo-complex was prepared by affinity purification followed by size exclusion chromatography (SEC), which showed that the Smc5/6 complex eluted in a single peak (Fig. 1A). The presence of all eight subunits in the peak fraction was confirmed by SDS-PAGE and by mass spectrometry analyses (Figs. 1B and S1B).

**Figure 1.**
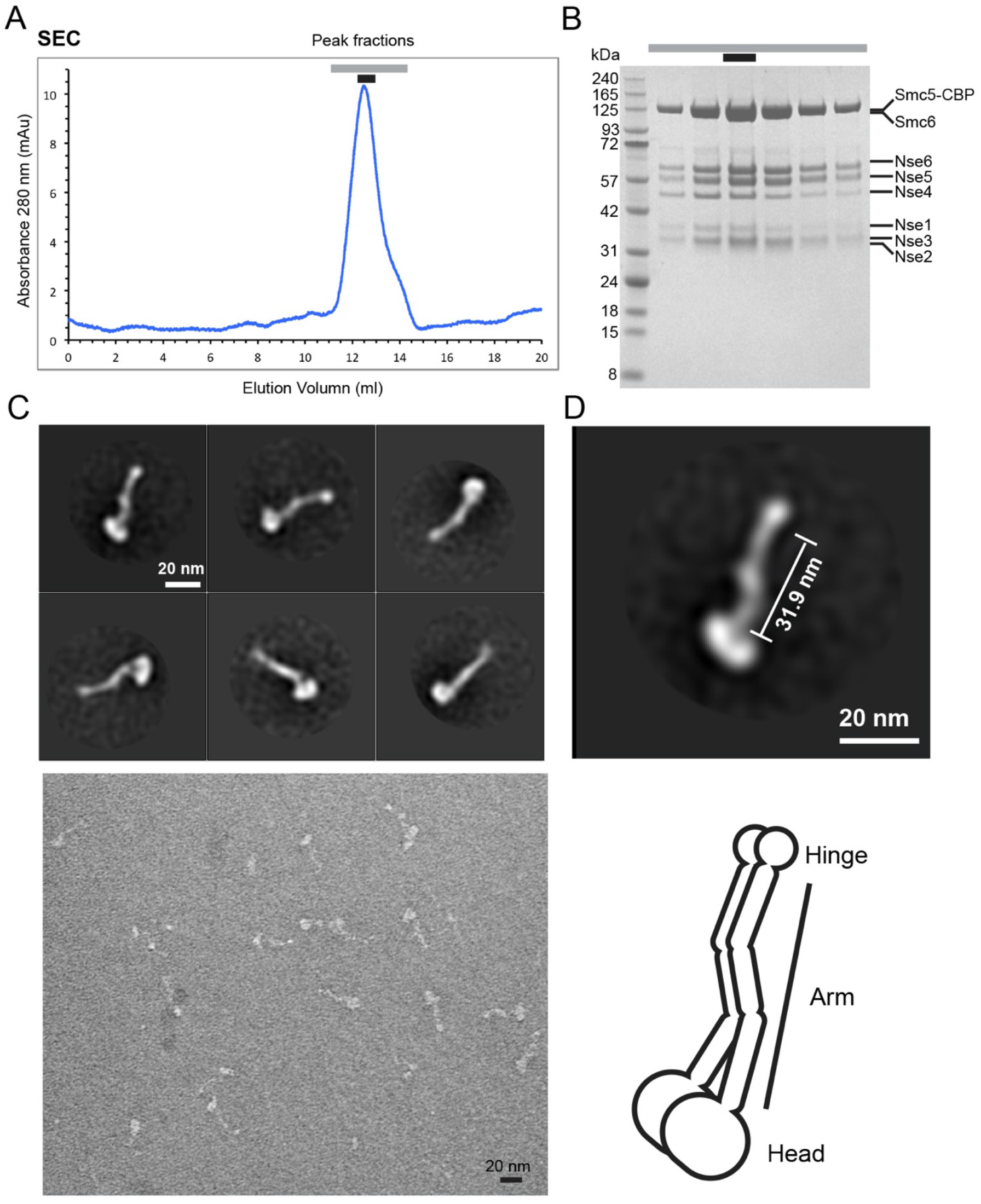
Negative stain EM analysis of the Smc5/6 holo-complex. (**A**) Size exclusion chromatography (SEC) profile of the Smc5/6 holo-complex. Peak fractions are indicated by lines above the elution profile from a Superose 6 Increase column. (**B**) Analysis of Smc5/6 holo-complex peak fractions by SDS-PAGE. A picture of the Coomassie stained gel is shown and the protein band corresponding to each subunit is indicated. The peak fraction marked by a black bar above the picture was used for negative stain EM analyses. (**C**) Negative stain EM analysis of the Smc5/6 holo-complex bound to ATPγS. Top panel: representative 2D class averages using a circular mask of 700 Å. Bottom panel: a typical field view of the negative stain EM of the Smc5/6 holo-complex. (**D**) 2D class averages of the Smc5/6 holo-complex with arm length labeled (top panel) and a cartoon of the overall structure of the complex (bottom panel).

Since ATP-binding affects SMC structures and the ATP-bound state is critical for SMC functions, we set out to assess the Smc5/6 holo-complex bound with ATPγS, a non-hydrolysable ATP analog (2–4). Negative stain EM data revealed that most particles of the Smc5/6 complex formed tripartite filaments, containing hinge, arm and head regions as seen for other SMC complexes (Fig. 1C). The complex in its ATPγS-bound state did not adopt an open-ring shape as previously speculated; rather, Smc5 and Smc6 appeared to align with each other throughout most of their length. Class averages generated by reference-free, two-dimensional (2D) image classification suggested that the Smc5/6 holo-complex contained a two-lobed head region, a small lobed hinge region, and an essentially linearly aligned rod-shaped arm connecting the head and hinge regions (Fig. 1D).

The EM analyses suggest that the arm region of the Smc5/6 complex is rather straight, with a notable thickening in the middle, likely stemming from Nse2 binding to the middle of Smc5 coiled-coil region as shown previously (Fig. 1D) (25). An elbow-bent feature seen for cohesin and condensin, wherein roughly one third of the coiled-coil regions proximal to the hinge bends back (Fig. S1A)(5–8) was not seen for holo-Smc5/6 in the presence of ATPγS. To further assess the arm structure, we compared the arm length of the Smc5/6 complex with those of cohesin and condensin. We measured the Smc5/6 complex arm length to be approximately 31.9 nm (Fig. 1D). This is 25% to 34% longer than the arm length of elbow-bent yeast condensin (25.5 nm) and cohesin (24.0 nm), respectively (5, 8). Because the coiled-coil sequences of Smc5 and Smc6 are about three-quarters in length of those of cohesin and condensin (8), an elbow-bent structure of Smc5/6 would predict an arm length shorter than those of cohesin and condensin, which was not the case. These analyses suggest that the arm region of ATPγS-bound Smc5/6 complex does not adopt an elbow-bent configuration.

### CL-MS suggests non-bending, rod-like shape of the Smc5/6 complex

Next we applied cross-linking mass spectrometry (CL-MS) to determine the topology of the Smc5/6 complex in solution. Purified Smc5/6 holo-complex bound with ATPγS was crosslinked with DSSO (disuccinimidyl sulfoxide) and CDI (N,N’-Carbonyldiimidazole) and analyzed by liquid chromatography nanoelectrospray ionization high-resolution mass spectrometry in three biological replicates (Fig. S1C). In total, we observed 337 unique cross-linked sites involving all eight subunits (Fig. S1D; Table S1). Among these, 214 were intra-molecular and 123 were inter-molecular crosslinks. We examined these results alongside the only known structure of the budding yeast Smc5/6 complex that includes Nse2 and its associated Smc5 coiled-coil region (25). Mapping all twelve unique amino acid cross-links involving these regions onto the Nse2-Smc5 structure showed that they were within the expected distance constraints for each cross-linker, less than 30 Å and 20 Å for DSSO and CDI, respectively (Fig. S1E). This is consistent with the accuracy of CL-MS observed in prior studies (26), and indicates high confidence in detected interactions.

A substantial fraction of the detected cross-links involved Smc5 and Smc6, with 87 intra-Smc5, 80 intra-Smc6, and 44 Smc5-Smc6 inter-molecular cross-links (Table S1). These crosslinks spanned most of the Smc5 and 6 sequences, with an enrichment in their arm regions (Figs. 2A, 2B, and S1D). The abundant Smc5 and 6 cross-links allowed us to derive features of their topology. First and as expected from antiparallel folding of both proteins, numerous intra-Smc5 and intra-Smc6 cross-links showed association of their N- and C-terminal coiled-coil segments (Fig. 2A). Second, twenty-eight cross-links connected the coiled-coil regions of Smc5 with those of Smc6 in a manner consistent with a zipped-up arm configuration (Fig. 2B). The apparent absence of cross-links between some parts of the arm regions may suggest local separation (Fig. 2B). Third, the two hinge regions formed multiple inter-molecular cross-links, consistent with their dimerization (Fig. 2B) (20, 24). The fewer intra-or inter-head cross-links suggest that the two heads may be shielded from cross-linking due to association with multiple Nse subunits (Figs. 2A, 2B, and S1D; Table S1). Overall, our CL-MS data are in agreement with the negative stain EM data and provide greater resolution on the manner of Smc5 and 6 association within the complex.

**Figure 2.**
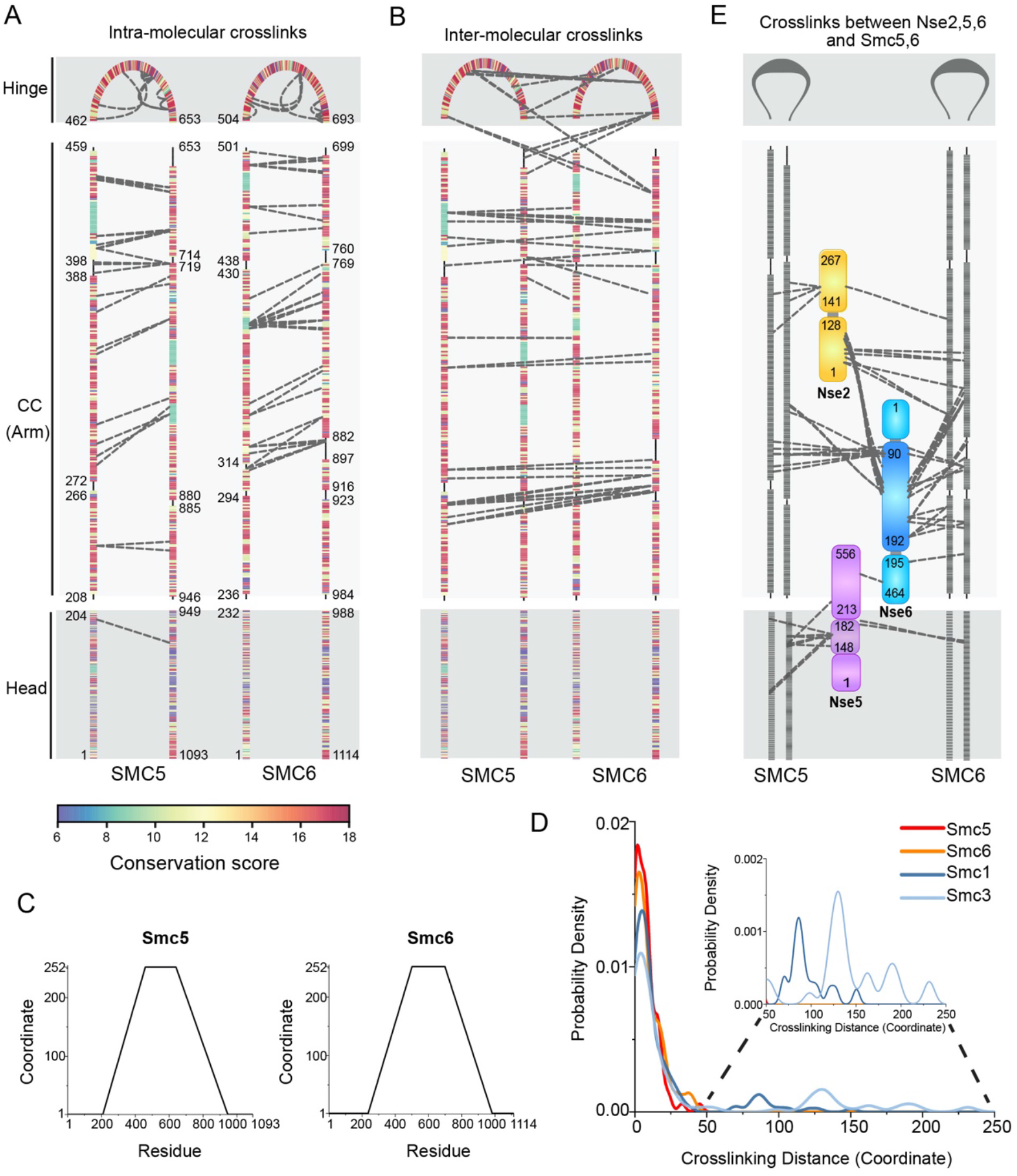
Cross-linking mass spectrometry analysis of Smc5/6 holo-complex. (**A**) Intra-Smc5 and intra-Smc6 cross-links. Three Smc5 and Smc6 regions, namely head, coiled-coil (CC, or arm), and hinge, are marked and residues bordering each region are labeled. Crosslinks connecting the N- and C-terminal portions of each region are represented by dashed lines between the corresponding amino acid pairs. Cross-links connecting adjacent sequences listed in Table S1 are omitted to highlight the antiparallel nature of the coil-coiled pairing of Smc5 and Smc6. Coiled-coil regions are interrupted by non-coiled-coil segments and residues bordering each predicted coiled-coil segment are indicated. Amino acids are colored based on conservation scores as shown in Fig. S1F and Table S3. (**B**) Inter-molecular cross-links between Smc5 and Smc6. The graph is presented as in panel A. (**C**) Transformed amino acid coordinates for Smc5 and Smc6 (detail see Methods). (**D**) Probability density of cross-linking distance based on transformed coordinates for Smc5 and Smc6 in the Smc5/6 complex, and for Smc1 and Smc3 in cohesin. CL-MS data for Smc1 and 3 are based on Burmann et al (8) (detail see Methods and Table S2). **(E)** Inter-subunits cross-links amongst Nse2, Nse5 and Nse6, and between them and Smc5/6. Smc5 and Smc6 are presented similarly as in panels A and B. Domain structures of Nse subunits are not drawn to scale to highlight the crosslinks.

Cross-linking distance analyses of cohesin have shown that 4-20% of Smc1 and Smc3 cross-links at their coiled-coil regions are separated by more than 100 amino acids (8). Such longdistance cross-links are consistent with extensive contacts between hinge-and head-proximal coiled-coil regions of cohesin caused by elbow bending (8). In contrast, all intra-protein crosslinks that fall onto the coiled-coil regions of Smc5 and Smc6 showed a distance of less than 45 amino acids (Figs. 2C and 2D; Table S2). In addition, no head-hinge cross-links were observed for holo-Smc5/6 (Table S1). Both of these observations are incompatible with a bent elbow model; rather they support the conclusion from our EM analyses that ATPγS-bound holo-Smc5/6 adopts a straight, rod-shaped configuration.

### Association of unique Nse subunits with Smc5/6 arm regions

Among the six Nse subunits, Nse2 and Nse5-6 are unique to the Smc5/6 complex, whereas Nse1 and Nse3 resemble the KITE subunits of bacterial SMC complexes, and Nse4 is similar to the kleisin subunit seen in most SMC complexes (13-15, 27). Nse1, Nse3, and Nse4 form a subcomplex (Nse1-3-4) that binds to the Smc5/6 head regions (20, 28). Accordingly, we found that Nse1, Nse3 and Nse4 cross-linked near the Smc5-6 head regions (Fig. S1D; Table S1). In contrast, Nse2 and Nse5-6 showed numerous cross-links with Smc5 and 6 arm regions (Figs. 2E and S1D; Table S1). Consistent with earlier structural data, Nse2 cross-linked to the middle section of the coiled-coils of Smc5 and the corresponding coiled-coils of Smc6 (Fig. 2E) (25).

Interestingly, a region spanning one hundred amino acids at the Nse6 N-terminus (90 to 192 a. a.) showed multiple crosslinks with Smc5 and Smc6 arm regions adjacent to the Nse2-binding site (Fig. 2E). We refer to this Nse6 region as the coiled-coil associated or proximal domain (CAD). Consistent with the observation that Nse2 and Nse6-CAD cross-linked to adjacent arm regions, cross-links between them were also observed (Fig. 2E). This suggests that the unique Nse2 and Nse6 subunits are also physically close to each other within the Smc5/6 complex.

Association of SMC mid-arm regions with non-SMC subunits has not been seen in other SMCs (2–4), indicating a highly specific feature of Smc5/6. In line with this conclusion, the arm sequences of Smc5 and Smc6, unlike those in Smc1/3 (cohesin) and Smc2/4 (condensin), are conserved (Figs. 2A and S1F) (9). In fact, conservation scores of Smc5 and Smc6 coiled-coil sequences are comparable to or higher than those of their head and hinge sequences (Fig. S1F; Table S3). This is consistent with the observation that the Smc5/6 arm regions uniquely engage in Nse interactions. Overall our analyses suggest that the Smc5/6 complex arm sequences are distinctively conserved and associate with Nse subunits not found in other SMC complexes. In principle, these interactions could limit the flexibility of the arm regions of the complex.

While the N-terminal CAD of Nse6 is in close proximity with Smc5/6 arm regions, its C-terminal region cross-linked with Nse5, consistent with them forming a subcomplex (Fig. 2E) (20, 29). However, unlike Nse6, Nse5 cross-linked to the Smc5 and Smc6 head regions (Fig. 2E). The Nse5 residues involved in these linkages were mainly restricted to a small segment in the middle of the protein (148 to 182 a. a.), referred to as the head associated or proximal domain (HAD) hereafter. In line with Nse5 being close to the Smc5 and Smc6 head regions, cross-links were found between Nse5 and the Nse1-3-4 subcomplex, which is also situated close to the head regions (Fig. S1D; Table S1). We did not observe crosslinks between Nse1-3-4 and the more hinge-proximally located Nse2 and Nse6 subunits (Fig. 2E; Table S1). In summary, our CL-MS data enable us to place the six Nse subunits relative to each other and to Smc5 and 6. Given that the SMC arm regions can affect the conformations and functions of the entire complex, the distinct features of the Smc5/6 arm regions, such as no elbow-fold and Nse association, can have important implications for its unique functions (see Discussion).

### Structure of the Nse5-6 complex reveals distinct domains used for an interaction network

Although the Nse5-6 subcomplex is one of the most unique features of the Smc5/6 complex, its structure has not been solved till now. Furthermore, it is a matter of debate whether Nse5 and Nse6, enriched in alpha helices, are similar to the HEAT repeat subunits of cohesin and condensin (1, 27). These HEAT repeat subunits, consisting of tandemly repeated modules of two alpha helices linked by a short loop, can form flexible structures and bind DNA (30). Both features are critical for cohesin and condensin entrapping and extruding DNA (30). As HEAT repeats are difficult to predict based on sequence analysis, we determined the Nse5-6 structure using cryo-EM.

Nse5 and Nse6 proteins were co-expressed and affinity purified. Their heterodimer eluted as a single peak in SEC (Fig. S2A), with the high purity of the complex confirmed by SDS-PAGE analyses (Fig. S2B). 2D class averages of cryo-EM data revealed the overall walnut shape of the complex (Fig. S2C). 3D classification, 3D auto-refine, and subsequent gold-standard refinement procedure (Figs. S2C and S2D) resulted in a reconstruction with an overall resolution of 3.2 Å (Fig. S2D and S2E; Table S4). Analyzing local resolution indicated that the Nse5 C-terminal domain (CTD; 445-556 a. a.) contributed to low resolution density (Figs. S2F and 3A). In addition, the N-terminal region of Nse6 (1-194 a. a.) was invisible in the density map, presumably due to structural flexibility. This region of Nse6 contains both the CAD element identified in this study for coiled-coil association and the Rtt107-interacting motif (RIM) reported previously (Fig. 3A) (22).

**Figure 3.**
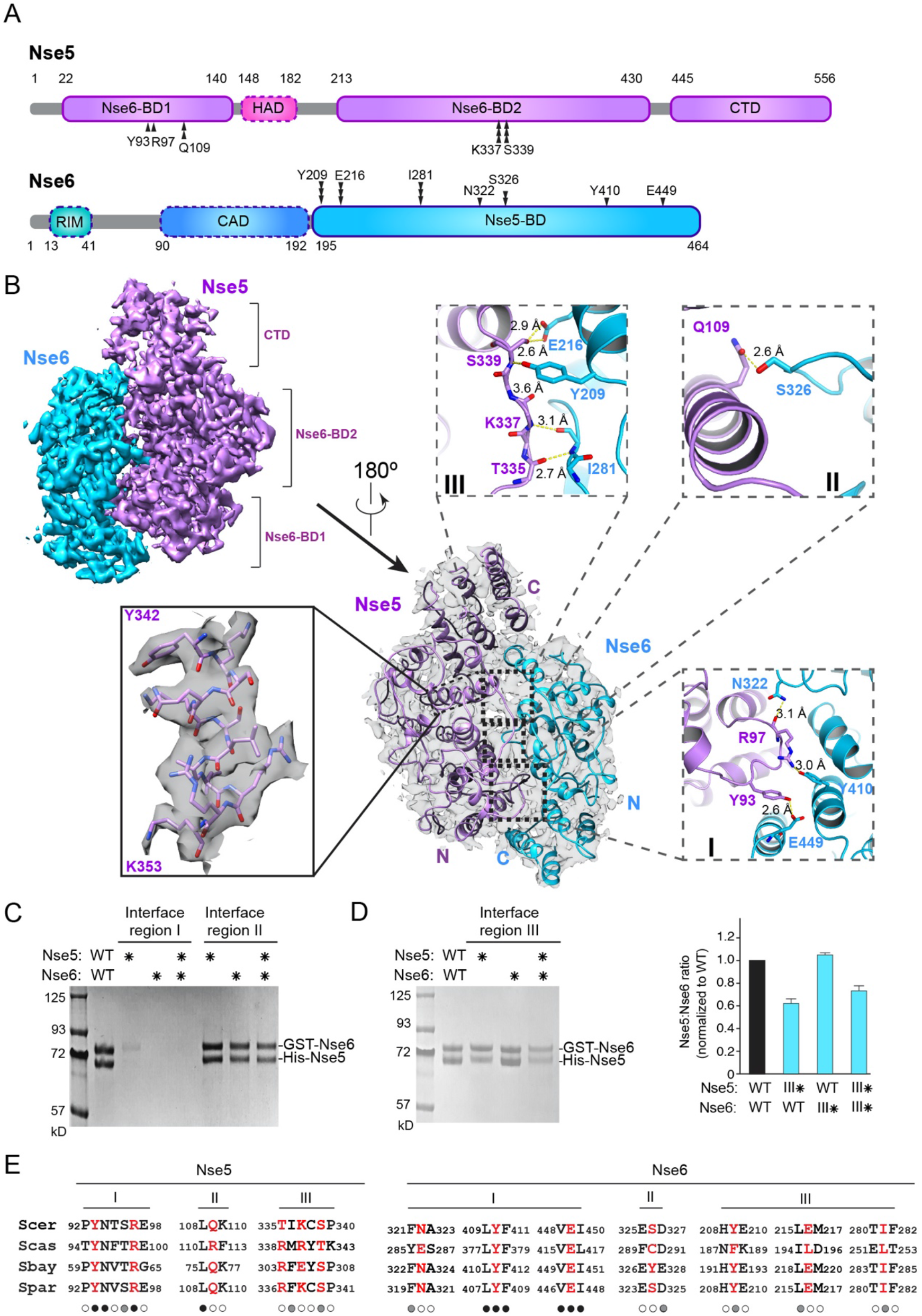
Cryo-EM structure of the Nse5-Nse6 complex. (**A**) Domain organization of Nse5 and Nse6 proteins. Residues involved in Nse5 and Nse6 interaction at regions I, II and III are marked by single, double and triple vertical arrows, respectively. The domains seen in the cryo-EM structure are outlined in solid lines while dotted outlines indicate the other domains. (**B**) Cryo-EM structure of the Nse5-6 complex. Two views of the structure are shown with 180° rotation, one with a transparent surface containing ribbon model (center) and one with surface rendering (upper left). An example of secondary structure elements with amino acid side chains is shown (left). Inlets depict detailed interactions between Nse5 and Nse6 at their binding regions I, II and III. Yellow dashed lines indicate the distances between atoms. (**C**) Mutating Nse5 and Nse6 interaction regions I and II has different effects on the Nse5-Nse6 complex levels. A representative SDS-PAGE gel picture of *in vitro* two-step pull-down results for wild-type or mutant Nse5 and Nse6 proteins. Asterisks indicate mutated form of the proteins. Region I mutations include Nse5-Y93, R97A and Nse6-N322, Y410, E449A. Region II mutations include Nse5-Q109A and Nse6-S326W. (**D**) Mutating Nse5 residues at the interaction region III reduces its binding to Nse6. Data are presented as in panel C, except Nse5-K337, S339W and Nse6-Y209A, I281W mutant proteins were examined. The graph on the right depicts the relative ratios of Nse5:Nse6 in the pull-down assays based on quantification from two experiments with average and standard deviations shown. (**E**) Sequence alignment of Nse5 and Nse6 in their interaction region I, II and III. Four *Saccharomyces* species were examined. *Scer: Saccharomyces cerevisiae, Scas: Saccharomyces castellii, Sbay: Saccharomyces bayanus, Spar: Saccharomyces paradoxus*. Residues involved in Nse5 and Nse6 interaction are colored red. Identical and similar amino acids are indicated by black and grey circles, respectively, while nonconserved amino acids are indicated by open circles below the sequences.

The resolution of the core of the Nse5-6 complex was approximately 3.0 Å (Fig. S2F) and hence we were able to build a *de novo* atomic model (Fig. 3B). Based on this model, Nse5 and Nse6 associate with each other largely in an antiparallel fashion (Fig. 3B). Nse6 uses its C-terminal half, referred to as Nse5-binding domain (Nse5-BD, 195-464 a. a.), to bind to two regions of Nse5: the Nse6-binding domain 1 (Nse6-BD1) located at its N-terminus (22-140 a. a.) and domain 2 (Nse6-BD2) that is more C-terminally situated (213-430 a. a.) (Fig. 3A). These two domains of Nse5 flank the HAD domain associating with the head (Fig. 3A). In summary, our data suggest that Nse5 and Nse6 use several distinct domains to engage in a network of interactions, including binding each other, anchoring to arm regions, contacting the head regions, and binding to Rtt107.

### Nse5 and Nse6 structural and functional features

Our structure shows that Nse5 and Nse6 are made up of mainly α-helices interrupted by loops and are devoid of β-sheet elements; however, these helices do not form HEAT repeat units (Fig. 3B). Consistent with this observation, while the HEAT repeat subunits in cohesin and condensin bind to DNA, Nse5-6 did not interact with either single stranded (ss) or double stranded (ds) DNA (Figs. S3A and S3B). As a comparison, we also examined the Nse1-3-4 subcomplex, as the homologous complex in human and fission yeast Smc5/6 complex associated with DNA (31). Indeed, we found that purified budding yeast Nse1-3-4 subcomplex interacted with both ssDNA and dsDNA (Figs. S3A and S3B), suggesting that Nse1-3-4 DNA binding is a highly conserved feature and is likely mainly responsible for Nse-mediated DNA binding in the Smc5/6 complex. Our DALI search (32) and structural alignment analysis showed no homologous structures related to Nse5, Nse6, or their complex in the PDB (protein data bank) to date, suggesting that Nse5-6 adopts a novel structure.

The dimeric interface of Nse5 and Nse6 is extensive, covering 1,942 Å^2^ area and consisting of three regions (Fig. 3B). Region I is the largest amongst the three, with side chains and main chains of two Nse5 residues (Tyr93 and Arg97) forming multiple polar interactions with side chains of three Nse6 residues (Asn322, Tyr410 and Glu449) (Fig. 3B). Region II is the least extensive, with a single side chain from Gln109 of Nse5 contacting the side chain of Ser326 of Nse6 via polar interactions (Fig. 3B). Interaction region III showed intermediate levels of contact compared with regions I and II. Here, polar interactions were formed between the main chains of Thr335, Lys337 and Ser339 in Nse5 and main chain of Ile281 and side chain of Tyr209 in Nse6, as well as between the side chain of Ser339 in Nse5 and side chain of Glu216 in Nse6 (Fig. 3B).

We performed mutational analyses to validate our structure of Nse5 and Nse6, and to assess the importance of the aforementioned three interfacial regions. Key residues supporting interactions in each region were mutated, and Nse5 and Nse6 mutations were examined individually or in combination. Co-expressed His6-tagged Nse5 and GST-tagged Nse6 were examined for complex formation in a two-step *in vitro* pull-down experiment. To this end, His6-Nse5 was first enriched by binding to Ni-NTA beads and the eluate was then bound to glutathione beads to assess the Nse5-Nse6 complex level. Compared with the wild-type Nse5-6 complex recovered, mutants showed different effects on the levels of the complex (Figs. 3C, 3D, and S3C). Correlating with their different interfacial structural features, mutations affecting region I caused the strongest disruption of Nse5 and Nse6 interaction, whereas those at region II had no detectable effects (Fig. 3C). Mutating region III residues led to a moderate reduction in Nse5-6 complex formation (Fig. 3D). The different levels of importance among the three Nse5 and 6 interfacial regions were consistent with their different degrees of sequence conservation. While region I residues show the strongest conservation, those in regions II and III are less well conserved (Fig. 3E). These analyses suggest that Nse5 and Nse6 form a stable dimeric complex and region I is the most critical in supporting their binding.

As an alternative way to assess our structural model, we mapped CL-MS data pertaining to Nse5 and Nse6 onto the structure. All seven crosslinks (six intramolecular and one intermolecular) mapped to the well-defined parts of the Nse5-6 structure were within the expected distance constraints of the respective crosslinker (Fig. S3D), thus providing another validation of our structure. In summary, our data suggest that Nse5 and Nse6 are distinct from the HEAT repeat subunits of cohesin and condensin at both structural and functional levels, and that Nse5-6 engages in protein-protein rather than protein-DNA interactions.

### Integrative structure modeling of a five-subunit Smc5/6 complex

The structures of Nse5-6 from this study and Nse2-Smc5 from our previous work (25), in conjunction with our CL-MS data, made it feasible to compute structural models. We omitted Nse-1-3-4 due to insufficient data to determine their juxtaposition in the complex. We used an integrative approach (33–37) to generate a structure of the five-subunit complex composed of Smc5, 6, and Nse2, 5, and 6 (designated Smc5/6-Nse2/5/6; Fig. 4A). The structural model was determined by combining comparative models of Smc5/6 head and hinge regions, parametrically designed backbone models of their coiled-coil regions, the structures of Nse2 and Nse5-6, and our CL-MS data (Methods and Fig. S4A).

**Figure 4.**
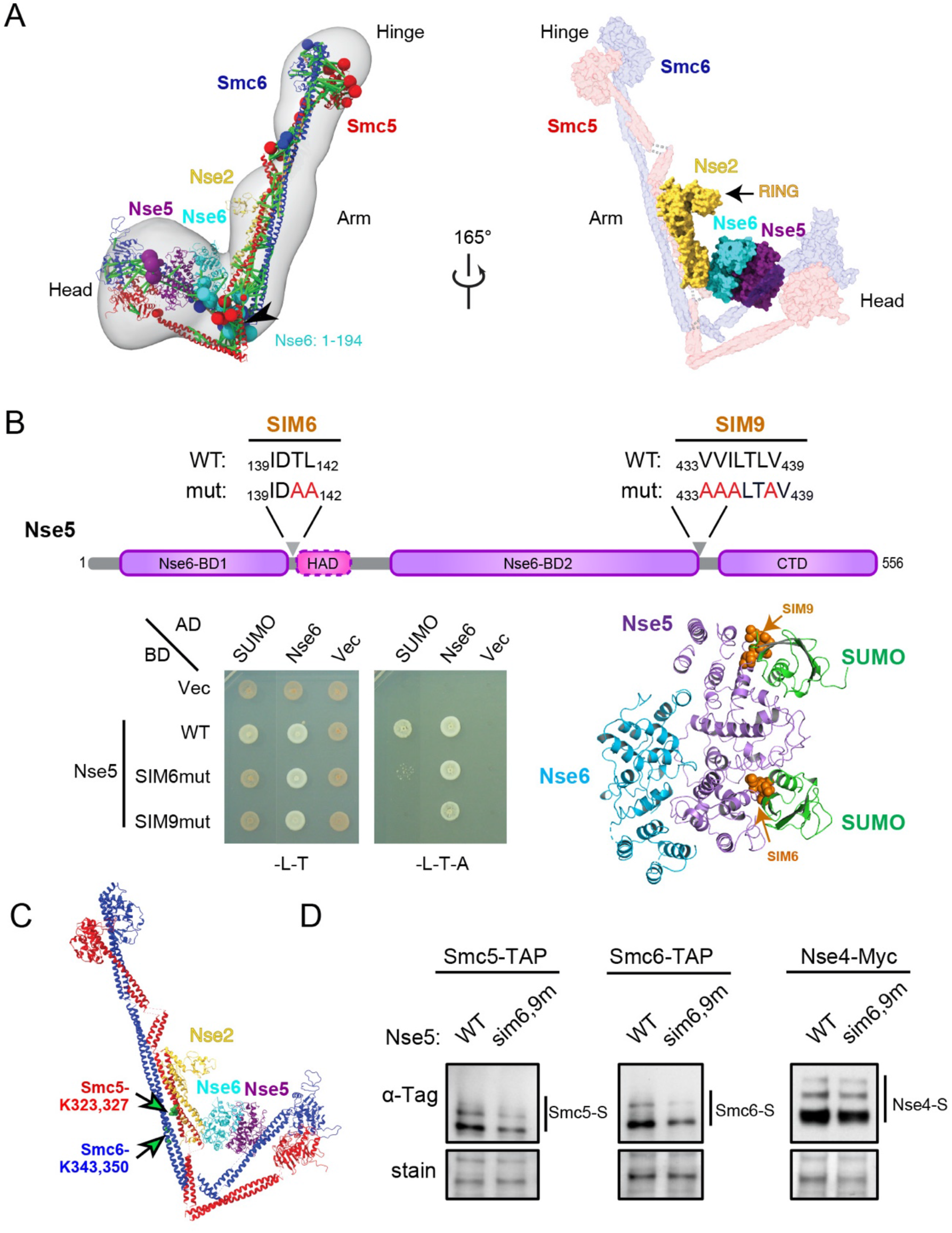
3D model of the Smc5/6 complex and Nse5 affecting Smc5 and 6 sumoylation. (**A**) Three-dimensional model of the five-subunit Smc5/6 complex. Left panel: Model representing the centroid of the ensemble of good scoring models obtained through Monte Carlo sampling. Regions with available atomistic structures were modelled as rigid bodies and shown in ribbon representation, while regions lacking atomistic structure were modelled as coarse-grained flexible beads and are shown with spheres whose radii are proportional to the average size of the fragment. The structural ensemble is demonstrated as a 3D localization probability density whose surface is rendered transparent for visual clarity. Satisfied crosslinks are shown in green while violated links are colored light brown. Right panel: surface presentation of the model highlights the proteins with available atomistic structure. (**B**) Nse5 SIM sequences contribute to its interaction with SUMO. Top: two SIM sequences are indicated in the Nse5 domain structure diagram. Bottom left: mutations in Nse5-SIM6 or SIM9 abolish the interactions with SUMO in yeast-two-hybrid assay. AD, Gal4 active domain; BD, Gal4 DNA binding domain; vec, vector. SC-Leu-Trp media (-L-T) selects for BD and AD plasmids, while SC-Leu-Trp-Ade media (L-T-A) reports for positive interactions. Bottom right: Nse5 SIM6 and SIM9 are located on the surface of the protein and accessible for SUMO binding based on structural modeling using the SUMO-SIM complex structure (PDB: 3V62,(52)). (**C**) Smc5 and Smc6 sumoylation sites in the 3D model of the Smc5/6-Nse2/5/6 complex. The Smc5 and 6 sumoylation sites, including Smc5 K323 and 327 and Smc6 K343 and 350 as mapped by Bhagwat et al. (44) are marked on our 3D model as green balls and indicated by arrows. (**D**) The sumoylation levels of Smc5 and Smc6, but not Nse4, were reduced in *nse5-sim6,9* mutant cells. Sumoylated proteins were enriched by Ni-NTA resins that bind to His8-tagged SUMO and examined by immunoblotting using antibodies recognizing the tag fused to the endogenous proteins. Loading is shown by Ponceau S stain (stain).

Figure 4A shows a representative model of the ensemble of structures obtained through structural sampling (see Supplemental Methods), for the five-subunit complex that sufficiently satisfies the input information. The structural ensemble has an excellent level of fit with our CL-MS data, with an overall 97.6% satisfaction (Fig. S4B). The structural precision of the ensemble, which refers to the average Cα RMSD of all models in the ensemble, from the representative centroid model, is 3.9 nm, and is twelve percent of the dimension of the Smc5/6 complex measured from the EM averages (31.9 nm; Figs. 4A and 1D). The model depicts an overall shape of the complex in agreement with our negative stain EM data. Specifically, Smc5 and 6 are paired at their hinge and head and aligned throughout coiled-coil regions without elbow bending (Fig. 4A). The middle section of the arms of the complex is thickened due to Nse2 binding (Fig. 4A). The model showed that the head-proximal coiled-coil regions, similar to the ‘neck’ region in cohesin and condensin, turned towards the hinge. We caution that while this configuration may suggest flexibility of these regions, it could change upon inclusion of the Nse1-3-4 subcomplex, which binds to these coiled-coils and the head regions. In the model, Nse5-6 is placed above the head region of the complex, with Nse6’s CAD (90-192 a. a.) in close proximity to Smc5 and Smc6 arm regions and Nse2 (Fig. 4A). The spatial closeness between Nse2 and Nse5-6 raises the possibility of their functional collaboration.

### Nse5 SUMO-interacting motifs affect Smc5 and 6 sumoylation

To test the possibility that the co-evolved Nse2 and Nse5-6 have functional connections, we considered the unique sumoylation function of the Smc5/6 complex that is not shared by other SMCs (16–19). Nse2 sumoylates Smc5, Smc6, and Nse4, among other substrates, and the sumoylation of Smc5 and 6 contributes to DNA replication and repair (16). However, the functional mechanisms of Nse2 as a SUMO ligase remain to be understood. Activities of other SUMO ligases require binding to the SUMO E2 through their RING domain and binding to the donor SUMO via SUMO interaction domains, and the latter interaction supports a productive conformation required for SUMO transfer (38). While Nse2 contains a RING domain, it has not been reported to interact with SUMO (25, 39), raising the possibility that Nse2 may require co-factors for optimal sumoylation efficiency for specific substrates.

Interestingly, we and an independent study found that Nse5 can associate with SUMO (40), raising the question whether it may assist Nse2-mediated sumoylation. To test this idea, we first located several four-amino acid sequences resembling SUMO-interacting motifs (SIMs) on Nse5 (Fig. S5A). SIMs harbor two to four hydrophobic residues and may contain an acidic residue (41–43). Taking advantage of our Nse5-6 structure, we assessed these sequences for accessibility for SUMO binding and found that only three SIMs (SIM 1, 6 and 9) fit this criterion (Fig. S5A). These and a few other SIMs were mutated and examined for SUMO and Nse6 association in yeast two hybrid assays (Fig. S5A). We found that mutating Nse5-SIM6 located between Nse6-BD1 and HAD, as well as SIM9 between Nse6-BD2 and CTD abolished SUMO interaction without affecting Nse6 association (Fig. 4B). In contrast, mutating SIM1 or other sequences reduced Nse6 interaction (Fig. S5B). We concluded that SIM6 and 9 of Nse5 primarily mediate SUMO interaction.

We proceeded to ask whether the SUMO binding of Nse5 contributes to Nse2-dependent sumoylation. Recently, mass spectrometry-based mapping showed that Smc5 and 6 are sumoylated at their coiled-coil regions (44). In our 3D model, these sites are located close to Nse2 and the Nse5-6 complex (Fig. 4C), thus are feasible to be influenced by both Nse2 and Nse5-6. Indeed, we found that the *nse5-sim6,9* mutant, which expressed at wild-type levels (Fig. S5C), was defective for the sumoylation of both Smc5 and Smc6 (Fig. 4D). This effect was specific because the sumoylation of Nse4 or several other Nse2 substrates was unaffected (Fig. S5D). Taken together, our *in vivo* data suggest that Nse5 SIMs favor the sumoylation state of the Smc5 and Smc6 proteins, providing a functional connection between the co-evolved Nse2 and Nse5-6 proteins.

## Discussion

The SMC family of proteins are primordial DNA modulators postulated to have evolved prior to histones (45). They are essential for maintaining the structures and functions of the genome. Extensive studies focusing on a subgroup of the SMC complexes, such as cohesin and condensin, have revealed their abilities to extrude DNA loops and cohese sister chromatids to sculpt chromatin (2–4). While the predominant view is that all SMC complexes behave similarly, multiple lines of evidence suggest that this may not apply to the Smc5/6 complex. Despite being essential for cell growth, the Smc5/6 complex does not appear to affect chromatin organization or topologically entrap chromatin (10–12). Rather, the Smc5/6 complex can directly regulate DNA replication and repair (13–15). The structural and molecular basis for the uniqueness of the Smc5/6 complex has been unclear. Our integrative, multi-scale structural and functional analyses of the yeast Smc5/6 complex have begun to address this knowledge gap and unveil several properties of this complex that are not shared by other SMC complexes.

Our negative stain EM, CL-MS and integrative modeling data corroborate each other and suggest that the arm regions of the Smc5/6 holo-complex in ATPγS bound form have four distinct and inter-related features. Our EM images and CL-MS analyses show linear alignment of the arm regions of Smc5 and Smc6, with thickening and small kinks in the middle, but no elbow bending as seen for cohesin and condensin. Our measurement of arm length from EM data and integrative modeling is consistent with the much-shortened arm sequences for Smc5/6 compared with cohesin and condensin as noted previously (8). Also, different from cohesin and condensin, Smc5/6 arm regions show interactions with its unique subunits, Nse2 and Nse6, which may provide a partial explanation for the lack of elbow bending. Further, Smc5/6 arm sequences show a high level of conservation, notably absent in cohesin and condensin. Altogether, these features allow us to propose that the arm regions of the Smc5/6 complex are engaged in multiple protein-protein interactions, and do not merely link the head and hinge regions. As the Smc5 arm region also binds to the Mph1 DNA helicase (46), it could serve as a platform for associating with additional genomic maintenance factors.

One of the most unique features of the Smc5/6 complex is its three co-evolved Nse subunits, namely Nse2 and the Nse5-6 complex. Our cryo-EM structure revealed a novel fold for Nse5-6 with a high α-helical content but devoid of HEAT repeats. Consistent with this, our biochemical experiments demonstrated Nse5-6’s lack of DNA binding ability, in contrast to Nse1-3-4. These features are a significant departure from the HEAT repeat subunits common to cohesin and condensin, which employ their DNA binding and flexibility to support multi-step DNA trapping and extrusion processes (30). The absence of these features in Nse5-6 suggests that the Smc5/6 complex is unlikely involved in chromatin organization in a way seen for cohesin and condensin. Rather, our CL-MS and cryo-EM data collectively showed that Nse5-6 contains distinct domains and engages in a network of protein-protein interactions. Besides the domains that form extensive interfaces in the Nse5-6 heterodimer, Nse5-HAD and Nse6-CAD are in close proximity with the head and arm regions, respectively. Recent yeast two-hybrid findings suggest that the human Nse6 homolog binds to the Smc5/6 arm regions, indicating that this interaction is evolutionarily conserved (47). Nse5-6 was reported to associate with the fragments of Smc5 and 6 hinge *in vitro* (20), but their crosslink in the holo-Smc5/6 was not seen here. While this interaction involving protein fragments may not reflect physiological association, we do not rule out other possibilities, such as the interaction may be involved in inter-complex association, and further clarification will be needed in the future. We have previously shown that Nse6 uses another domain for binding to Rtt107 that in turn recruits the Smc5/6 complex to damaged chromatin (22). Collectively, our data suggest that Nse5-6 builds a protein-protein interaction network that supports Smc5/6 complex function in genome maintenance.

Our integrative model points to a spatial proximity among the three co-evolved Nse subunits (Nse2. Nse5 and Nse6) unique to the Smc5/6 complex. Our *in vivo* data show that Nse5 contains SIM sequences that mediate its SUMO interaction. Further, Nse5-SIM mutations specifically reduce Nse2-dependent Smc5 and Smc6 sumoylation that occurs at their arm regions located close to Nse2 and Nse5-6. These data suggest one shared function between Nse2 and Nse5-6. We consider at least two non-mutually exclusive mechanisms for the roles of Nse5-SIM in Smc5 and 6 sumoylation. Given that other SUMO ligases binding to SUMO to support SUMO transfer to substrates, the apparent lack of SUMO binding by Nse2 suggests that Nse5 may substitute for it during specific sumoylation events. Interestingly, we recently found that the Esc2 protein acts as another cofactor that helps Nse2-mediated sumoylation of the STR complex and Pol2 (48). It is thus possible that Nse2 may collaborate with different cofactors to efficiently sumoylate distinct substrates, and that the Smc5/6 complex may act as a ‘composite SUMO ligase’. This model does not exclude another possibility wherein Nse5 simply helps to increase local SUMO concentration via binding to SUMO. Testing these and other potential models in the future will shed light into how the Smc5/6 complex acts as a SUMO ligase and uses sumoylation to regulate DNA transactions.

During the course of our work, negative stain EM and CL-MS studies of a human and a budding yeast Smc5/6 complex reached different conclusions, with only the latter suggesting an elbow-bent configuration (49, 50). This latter study (50) and our work have some technical and data interpretation differences and we additionally provided an integrative model that supports a straight-arm conformation of Smc5 and Smc6. However, our work examined the ATPγS-bound form of the Smc5/6 complex, in contrast to mixed populations of the complex used in the other studies (49, 50). This raises the possibility that Smc5/6 arm flexibility may be regulated by ATP. More work will be required to determine structural changes in Smc5/6 during its ATPase cycle and DNA interactions. These future studies will provide further insights into the observed DNA linking and sumoylation activities of the complex (16, 49-51).

Overall, our integrative structural analyses provide critical insights into multiple unique structural and functional features of the Smc5/6 complex. These include the straight Smc5/6 arm regions and its three unique and co-evolved Nse subunits that exhibit physical proximity and functional cooperation. We propose that these Nse proteins and their interacting arm regions can serve as platforms for binding and collaborating with DNA replication and repair factors, while Nse1-3-4 and other parts of Smc5 and 6 support DNA binding. Future work to elucidate how these two aspects of the complex collaborate to enable the Smc5/6 complex to carry out its described roles in DNA replication and repair will further our understanding of this multi-functional genome maintenance complex.

## Acknowledgements

We thank Shelly Lim and Jacob Bonner for assisting in yeast strain construction in the early stage of this work. We appreciate Prabha Sarangi, Shengliu Wang, and Dirk Remus for comments on the manuscript. Z.S. is supported by an A*STAR scholarship. This work is supported by the National Institute of General Medical Science grants R01GM080670 and R01GM131058 to X.Z., R01GM083960 and P41GM109824 to A.S., National Cancer Institute grant R01 CA214812 to A.K., Maloris Foundation to D.P., a Beene Cancer Center Grant jointly to X.Z. and D.P.. A.K. is a consultant for Novartis and Rgenta.

## Author contributions

Y.Y. carried out negative stain EM and cryo-EM studies under the supervision of D.P., and protein-protein and protein-DNA binding assays under the supervision of X.Z.. S.L. purified the Smc5/6 holo-complex, performed cross-linking assays, and conducted most *in vivo* studies, and K.C. examined Nse5-SIM mutants in yeast two-hybrid assays under the supervision of X.Z. Z.S. performed crosslink MS experiments and data analyses under the supervision of A.K. and X.Z. T.S. carried out integrative computational modeling under the supervision of A.S. and X.Z.. B.W. performed initial Nse5 and Nse6 protein characterization under the supervision of X.Z. All authors are involved in experimental designs and data analyses. Y.Y. S.L. Z.S, T.S. and X.Z. wrote the manuscript with input and edits from all the other authors.

## Materials and Methods

### Yeast strains and genetic methods and plasmids

All yeast strains are derivatives of W1588-4C, a *RAD5* derivative of W303 (*MATa ade2-1 can1-100 ura3-1 his3-11,15 leu2-3,112 trp1-1 rad5-535*). For other *in vivo* assays, at least two strains per genotype were examined in each experiment, and only one is listed in Table S5. Plasmids used are listed in Table S6. Detailed strain and mutation construction and standard yeast two-hybrid assay procedure are described in the Supplemental Materials and Methods.

### Purification of Smc5/6 holo-complex, Nse5-6, and Nse1-3-4 subcomplexes, and biochemical assays

The Smc5/6 holo-complex was purified from yeast cells while the two subcomplexes were purified in bacteria. Details of protein purification, *in vitro* pull down tests, and DNA binding assays are in Supplemental Materials and Methods.

### Negative stain EM of the Smc5/6 holo-complex and Cryo-EM analyses of the Nse5-6 complex

Purified SMC5/6 holo-complex were examined by a Philips CM10 electron microscope and images were recorded at a calibrated magnification of ×41513, yielding the pixel size of 2.7Å at specimen level. Purified Nse5-6 complex was applied onto glow-discharged UltrAuFoil and images were collected on a FEI Titan Krios electron microscope with a Gatan K3 camera with a 1.064 Å pixel size. Image processing was performed by RELION 3.0 and a total of 657,200 particles were selected for 3D classification. Details are in Supplemental Materials and Methods.

### Cross-linking mass spectrometry and data analyses

The Smc5/6 complex was cross-linked with DSSO (Thermo) or CDI (Sigma Aldrich), and purified before digestion with LysC followed by Trypsin. The mass spectrometry proteomics data have been deposited to the ProteomeXchange Consortium via the PRIDE (53) partner repository with the dataset identifier PXD023164. Details are in Supplemental Materials and Methods.

### Integrative structure determination of Smc5/6-Nse2/5/6 complex

Integrative structure modeling proceeded through the standard four-stage protocol (33–37), which was scripted using the *Python Modeling Interface* package, a library for modeling macromolecular complexes based on the *Integrative Modeling Platform* software (35), version develop-255ae6c (https://integrativemodeling.org). Details are in Supplemental Materials and Methods.

### Detection of *in vivo* protein sumoylation

For all proteins, except Pol2, examined here, SUMOylated proteins were pulled and probed for specific substrates by immune-blotting (54). For Pol2, the protein was immuno-precipitation before immune-blotting for its sumoylated forms. Detailed are in Supplemental Materials and Methods.

