## Supplemental Figure Legends for "Integrative analysis reveals unique features of the Smc5/6 complex"

**for**

**Supplemental Figure Legends**

**Figure S1. CL-MS data of the Smc5/6 complex and Smc5 and 6 sequence conservation**.

(**A**) Schematic showing typical SMC protein domain and fold.

(**B**) Protein sequence coverage of the subunits of the Smc5/6 holo-complex based on MS analyses. Bars are labeled with average sequence coverage (%) from 3 replicates cross-linked by DSSO and 3 replicates cross-linked by CDI. Error bars indicate standard deviation across all 6 replicates.

(**C**) A representative gel picture for examining the crosslinked ATPγS-bound Smc5/6 holo-complex.

(**D**) Circular plot showing crosslinks for ATPγS-bound Smc5/6 holo-complex by both DSSO and CDI crosslinker. Intra-protein cross-links are colored purple. Inter-protein cross-links are colored green.

(**E**) Mapping DSSO and CDI crosslinks to the Nse2-Smc5 structure (PDB ID: 3HTK, (35)) and histogram depicting Cα-Cα distances for mapped crosslinks.

(**F**) Plot of conservation score for Smc5 and Smc6 sequence with bin size of 10 a. a. Horizontal dotted line represents average conservation score across the whole protein sequence.

**Figure S2. Cryo-EM reconstruction of the Nse5-6 complex.**

(**A**) Size exclusion chromatogram (SEC) of Nse5-Nse6 complex. Peak fractions were labeled as line segments on the top.

(**B**) Analysis of Nse5-Nse6 complex peak fractions by SDS-PAGE. A representative Coomassie staining picture is shown. Band corresponding to each subunit is indicated.

(**C**) Workflow of cryo-EM image processing for the Nse5-Nse6 complex. Examples of cryo-EM 2D classification results of the Nse5-Nse6 complex are shown on the top.

(**D**) Global Fourier Shell Correlation (FSC) curve of the Nse5-Nse6 complex. The curve for the two half datasets is in blue and that for the refined model versus the cryo-EM map is in red. The overall cryo-EM map resolution is 3.2 Å with FSC set at 0.143.

(**E**) Angular distribution plot of final 3D EM map for the Nse5-Nse6 complex.

(**F**) Final 3D reconstructed map of the Nse5-Nse6 complex colored according to local resolution. The resolution of the majority of the map is 3 Å, with relatively poor density seen at the Nse5 C-terminal domain.

**Figure S3. Functional and structural features of the Nse5-6 complex.**

(**A**) Analysis of purified Nse5-6 and Nse1-3-4 complexes by SDS-PAGE. Representative Coomassie stained gel pictures are shown. Band corresponding to each subunit is indicated.

(**B**) *In vitro* DNA binding assay results. Fluorescein labeled single stranded (ss) and double stranded (ds) DNA were mixed with increasing levels of proteins at protein:DNA molar ratios from 1:1 to 1:8 as indicated. Reactions mixtures were analyzed on gel. Representative images of scanned fluorescent signals are shown.

(**C**) Examination of Nse5 and Nse6 proteins in the crude extract of bacterial cells induced for their expression. The presence of Nse5 and Nse6 proteins was detected by immunoblotting using anti-GST and anti-His antibodies. Labeling is as Figure 3C and 3D. Nse5 mutations affecting interface regions I and III supported protein expression, yet as shown in Figure 3C and 3D, were defective in the Nse5-6 complex formation. Nse6 mutations affecting interface regions I and III appeared to reduce Nse5 expression levels, however, this effect may not solely be responsible for the reduced Nse5-6 complex levels seen in Figure 3C and 3D, since similar level of Nse5 due to mutation in interface region II can support wild-type level of Nse5-6 complex formation.

(**D**) Mapped DSSO and CDI crosslinks in the structure of the Nse5-Nse6 complex, and histogram depicting Cα-Cα distances for the mapped crosslinks.

**Figure S4. Integrative model of the Smc5/6-Nse2/5/6 complex.**

(**A**) Four-stage scheme of integrative modeling, of the Smc5/6-Nse2/5/6 complex.

(**B**) Distribution of Cɑ-Cɑ distances of crosslinked residues for DSS (left subplot) and CDI (right subplot) with the threshold distances of 30 Å (for DSSO) and 20 Å (CDI) marked with dotted vertical lines. The overall crosslink satisfaction is 97.6%.

**Figure S5. Examination of Nse5’s SUMO binding motifs**

(**A**) Nse5 sites that match SIM consensus sequences. These sequences were assessed for their accessibility for SUMO binding and locations relative to the Nse5-Nse6 binding interface. Mutations made for testing SUMO binding are indicated in red.

(**B**) Summary of yeast two-hybrid results of Nse5 mutations in effecting SUMO binding.

(**C**) The *nse5-sim6,9* mutant does not affect its protein level.

(**D**) The effects of *nse5-sim6,9* mutation on sumoylation of non-Smc5/6 substrates. Nse2 substrates, including the Holliday junction dissolution complex, Sgs1-Top3-Rmi1, and the DNA polymerase Pol2, were maintained. Sgs1-Top3-Rmi1 sumoylation was examined as in Figure 4D. HA-tagged Pol2 was immunoprecipitated and its sumoylated form was detected using anti-SUMO antibody in immunoblotting, while unmodified Pol2 was detected using the anti-HA antibody.
