## Supplementary figures and images for "Integrative analysis reveals unique features of the Smc5/6 complex"

### Supplemental Figure 1

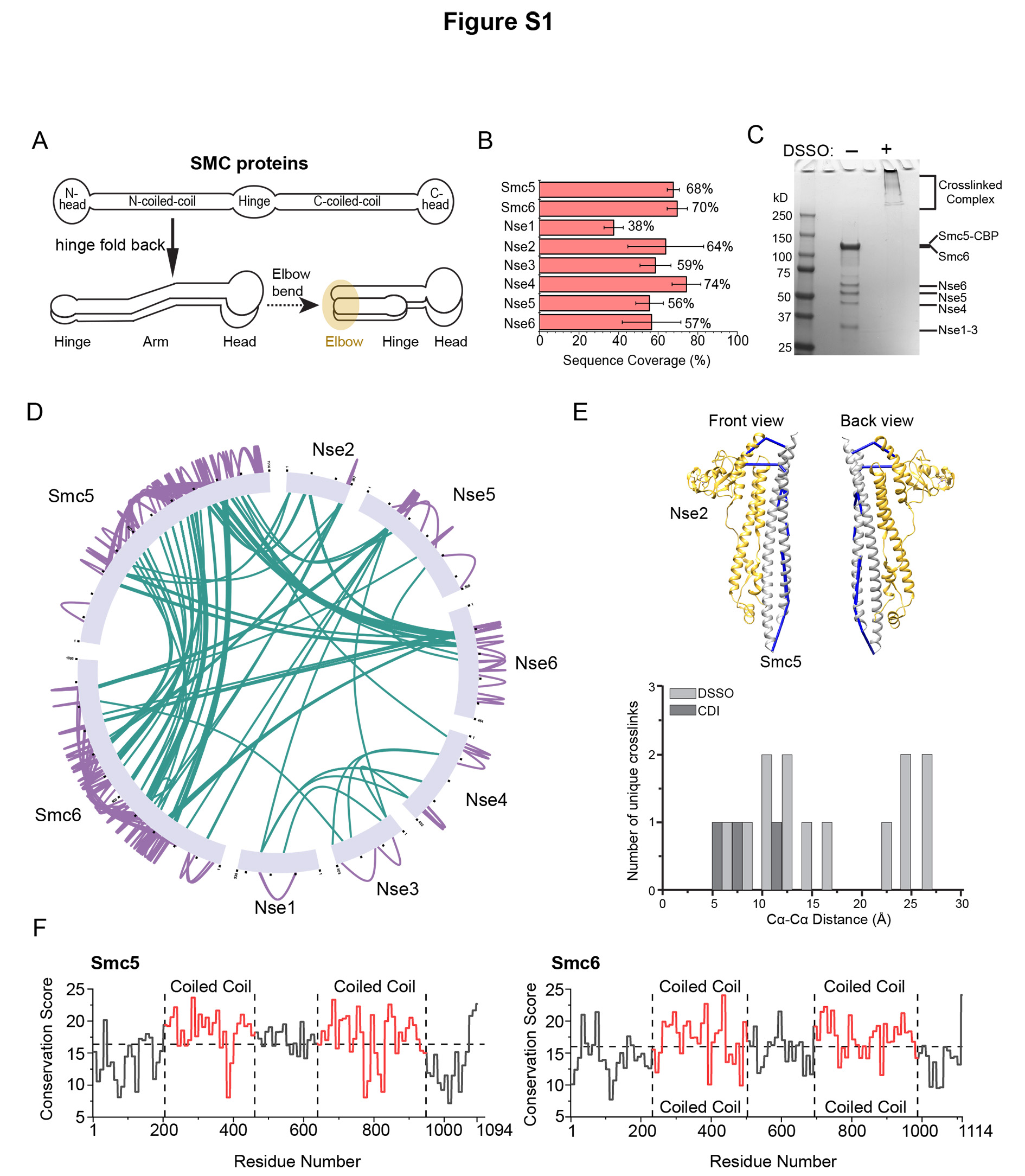

### Supplemental Figure 2

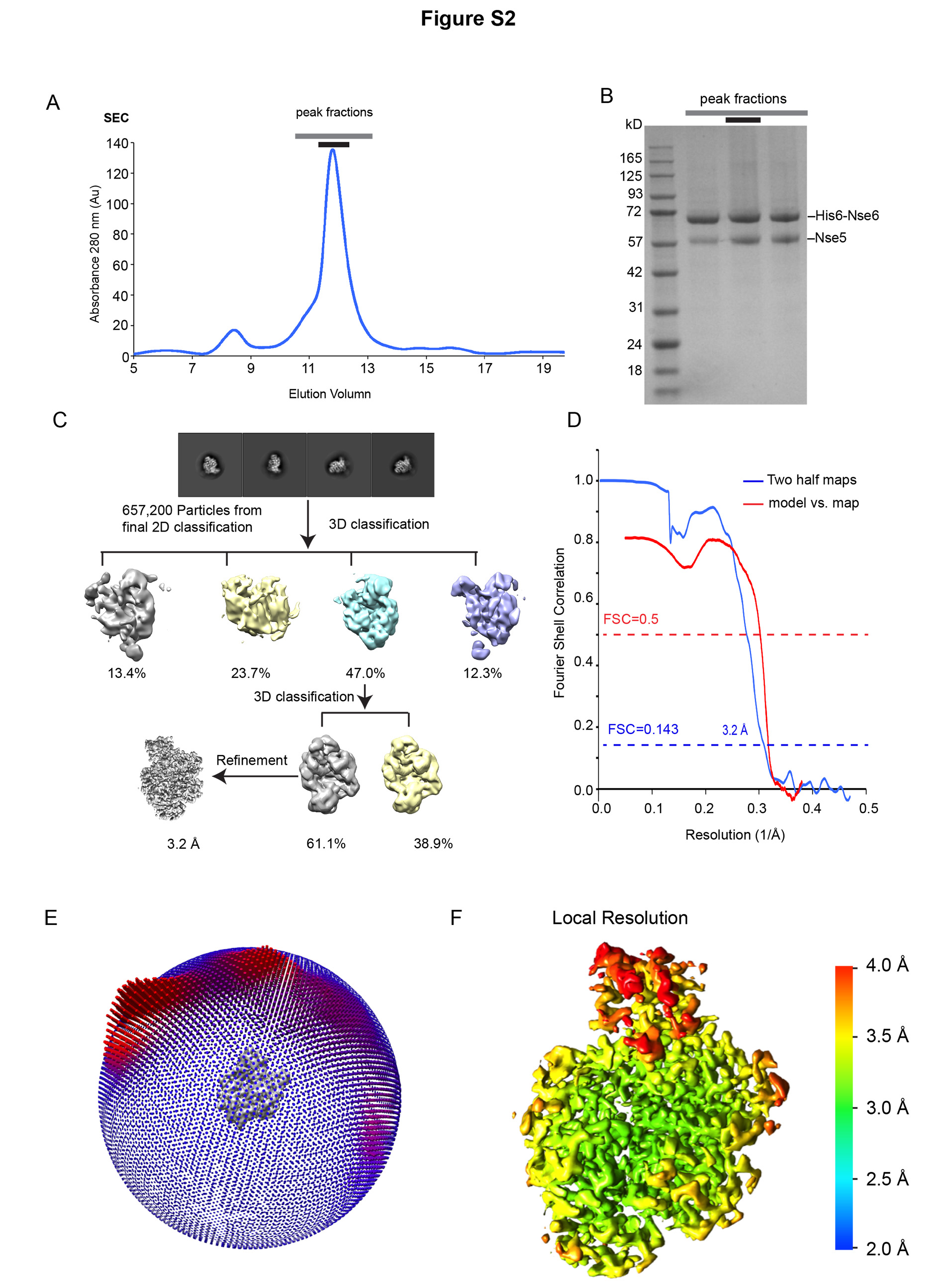

### Supplemental Figure 3

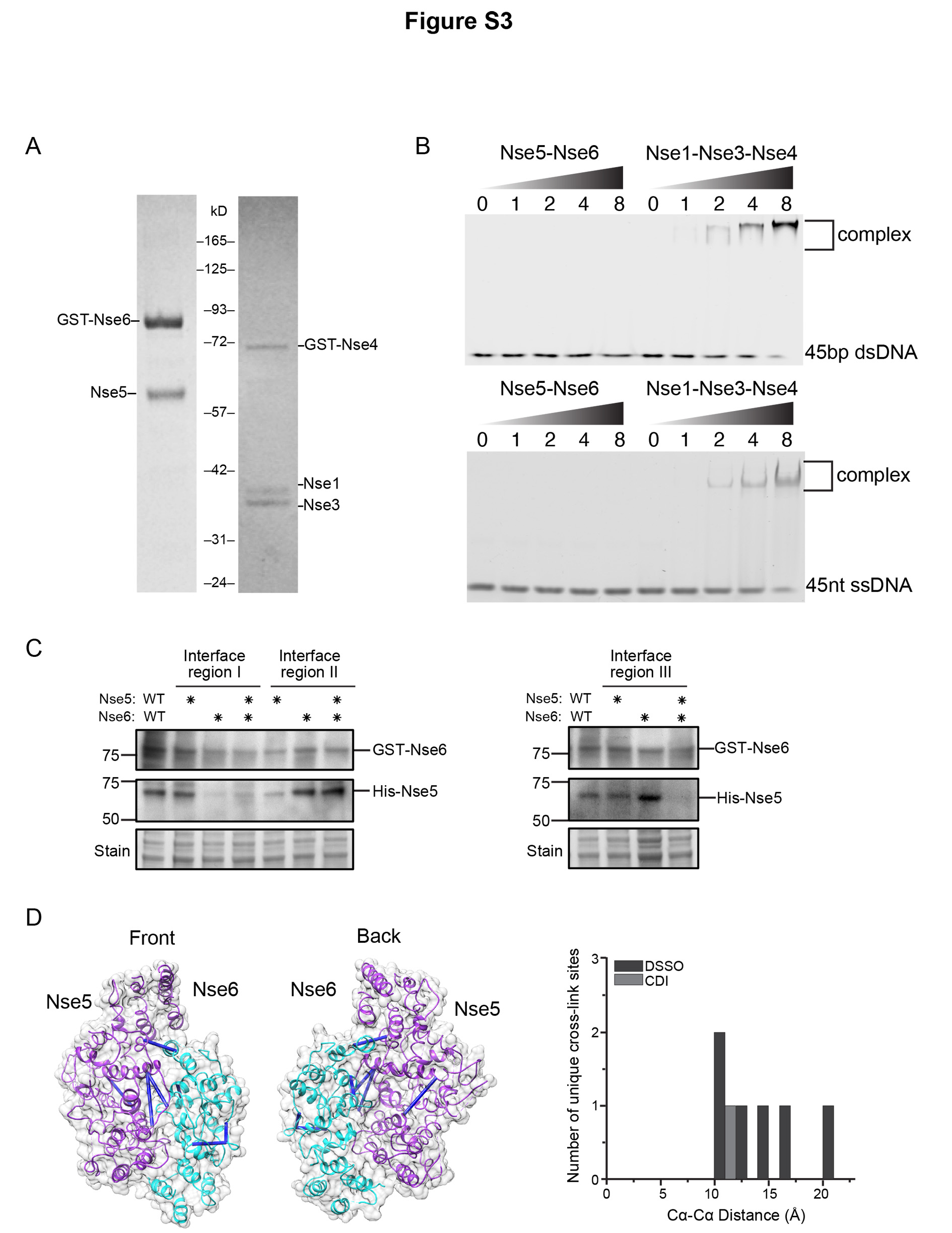

### Supplemental Figure 4

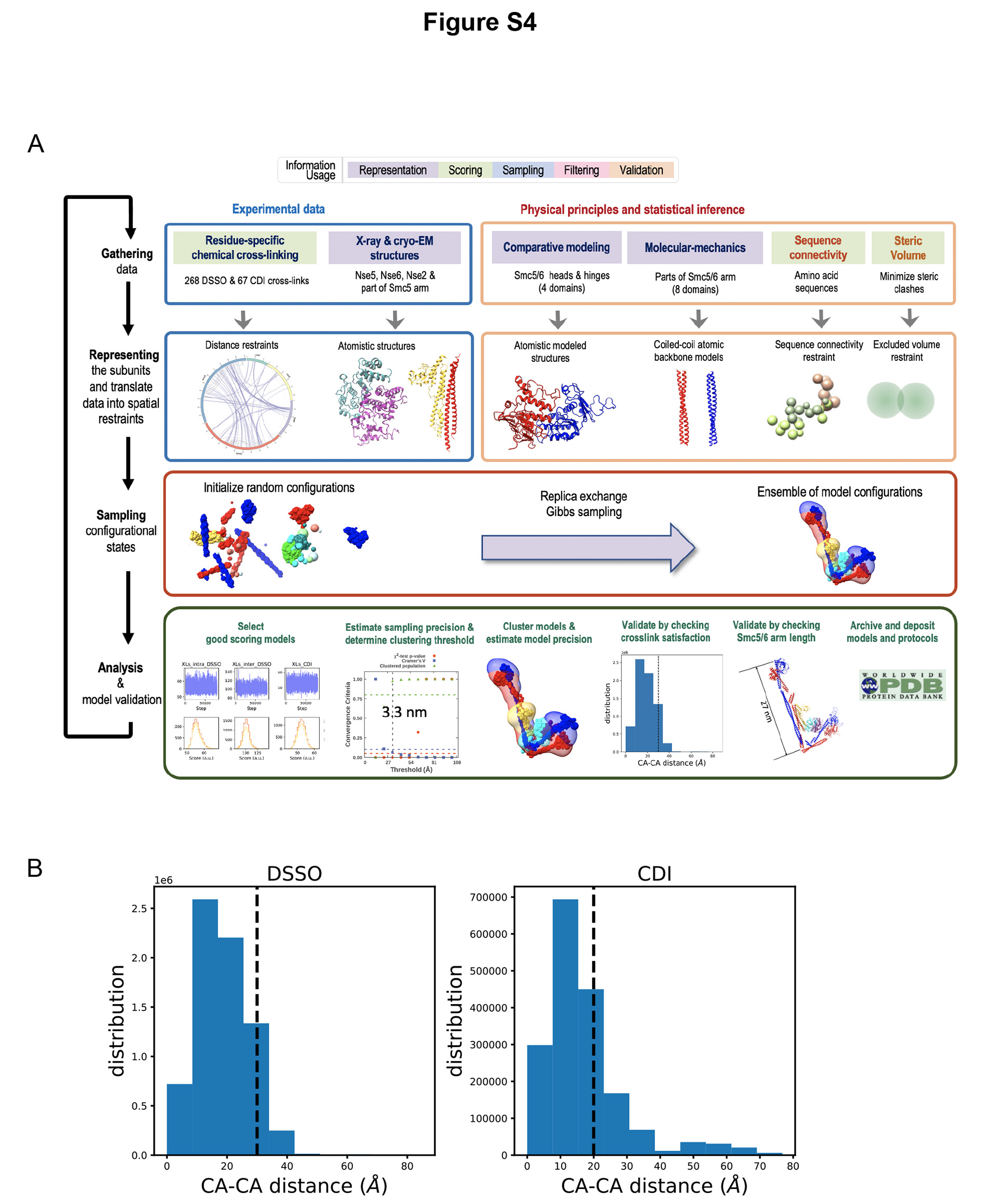

### Supplemental Figure 5

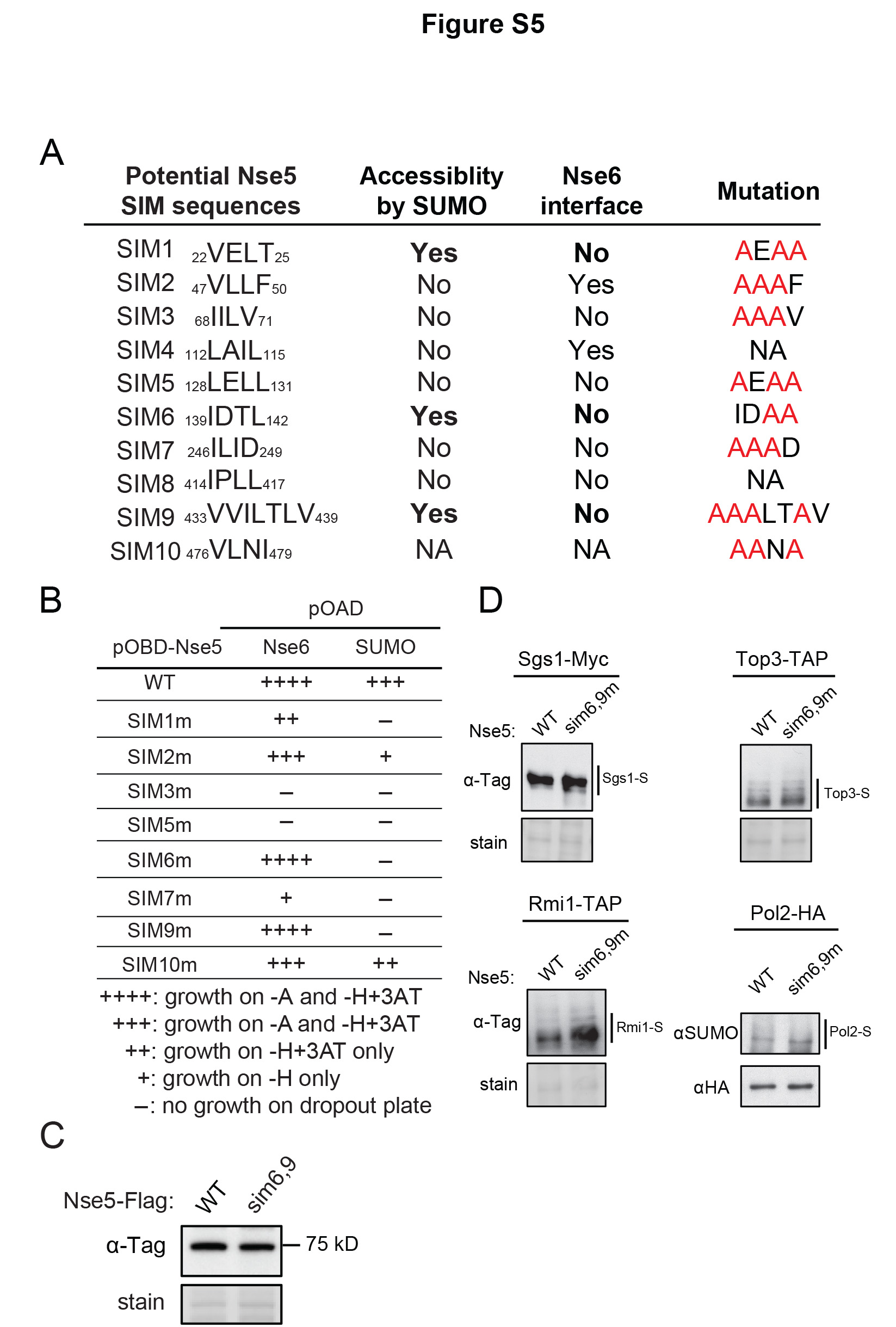
